## Supplementary 1 for "Caught in a trap: DNA contamination in tsetse xenomonitoring can lead to over-estimates of *Trypanosoma brucei* infection"

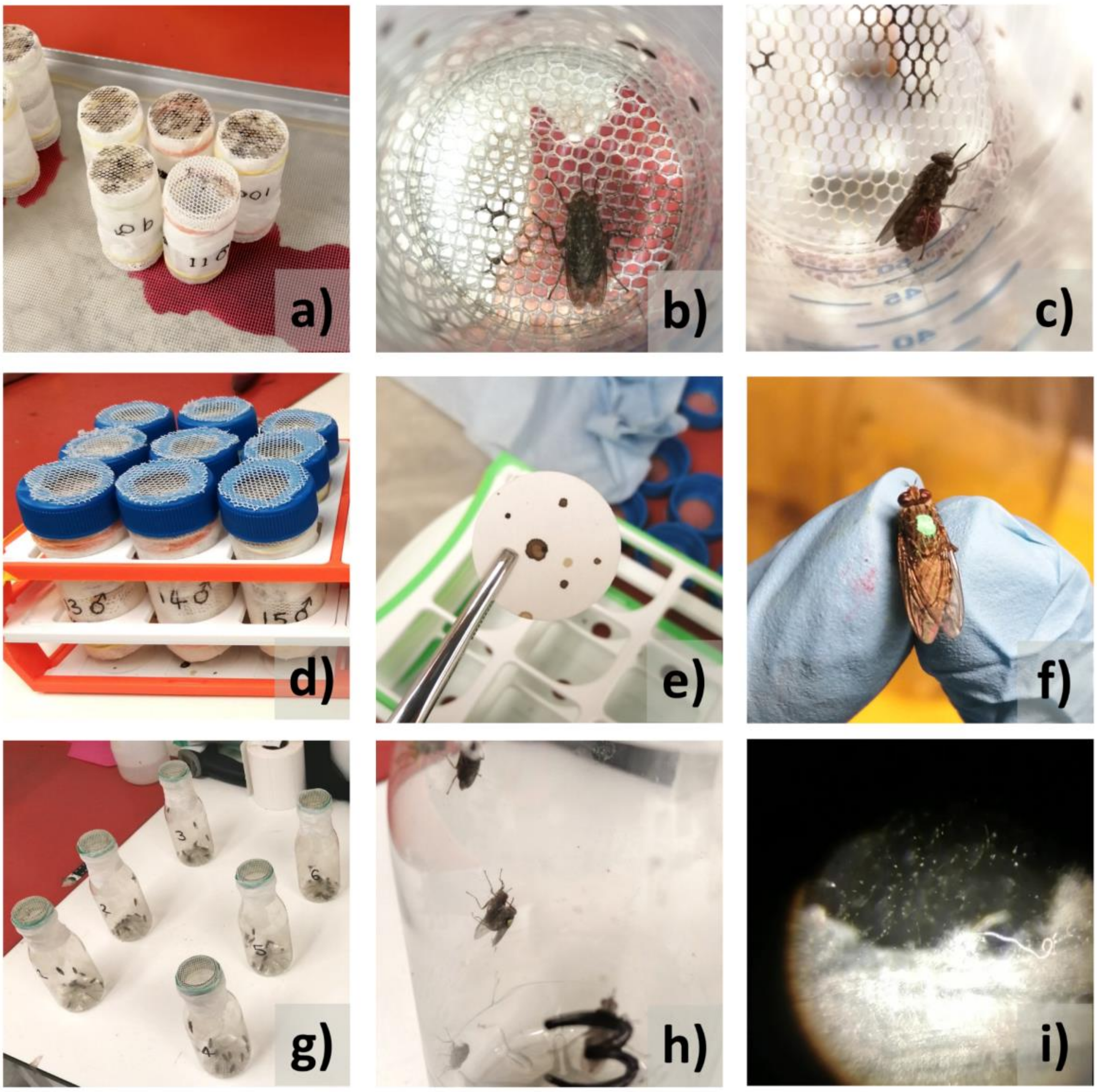


**S1: Images from experiments conducted on insectary-reared tsetse.** (a) and (b) show tsetse being blood-fed in solitary cells; (c) tsetse resting after bloodmeal; (d) tsetse solitary cells suspended above filter paper discs in rack; (e) collection of tsetse faecal samples on filter paper; (f) an infected tsetse marked with green oil paint; (g) experiment trap cages; (h) two infected tsetse (fly IDs 87 and 109) copulating inside trap cage during experiment; (i) dissected tsetse midgut infected with *T. brucei* as viewed under a microscope (400X).
