## Supplementary 2 for "Caught in a trap: DNA contamination in tsetse xenomonitoring can lead to over-estimates of *Trypanosoma brucei* infection"

| Treatment | Replicate | Infected | | | Uninfected | | |
|---|---|---|---|---|---|---|---|
| | | M | F | Total IF | M | F | Total UF |
| **T1** | **A** | 1 | 8 | 9 | 2 | 1 | 3 |
| | **B** | 4 | 5 | 9 | 2 | 1 | 3 |
| | **C** | 4 | 5 | 9 | 2 | 1 | 3 |
| **T2** | **A** | 4 | 2 | 6 | 2 | 4 | 6 |
| | **B** | 4 | 2 | 6 | 2 | 4 | 6 |
| | **C** | 4 | 2 | 6 | 2 | 4 | 6 |
| **T3** | **A** | 1 | 0 | 1 | 5 | 6 | 11 |
| | **B** | 1 | 0 | 1 | 5 | 6 | 11 |
| | **C** | 1 | 0 | 1 | 5 | 6 | 11 |
| **C0** | **A** | 0 | 0 | 0 | 5 | 7 | 12 |
| | **B** | 0 | 0 | 0 | 6 | 6 | 12 |
| | **C** | 0 | 0 | 0 | 6 | 6 | 12 |

**S2: A table displaying *G. m. morsitans* sex and infection ratios for trap cage experiments**. M = male, F = female, IF = infected fly, UF = naïve uninfected fly.
