## Supplementary 3 for "Caught in a trap: DNA contamination in tsetse xenomonitoring can lead to over-estimates of *Trypanosoma brucei* infection"

| Trap ID | Transect | Region | Latitude | Longitude |
|---|---|---|---|---|
| TA-1 | TA | Tarangire NP | 157791.8 | -432982 |
| TA-2 | TA | Tarangire NP | 157758.3 | -433093 |
| TA-3 | TA | Tarangire NP | 160220.6 | -433927 |
| TA-4 | TA | Tarangire NP | 160390.8 | -434067 |
| TA-5 | TA | Tarangire NP | 162344 | -435548 |
| TA-6 | TA | Tarangire NP | 162421.2 | -435634 |
| TA-7 | TA | Tarangire NP | 166310.1 | -439209 |
| TA-8 | TA | Tarangire NP | 166425.8 | -439176 |
| TA-9 | TA | Tarangire NP | 169880.1 | -440627 |
| TA-10 | TA | Tarangire NP | 169987.6 | -440676 |
| TA-11 | TA | Tarangire NP | 175093 | -440977 |
| TA-12 | TA | Tarangire NP | 175332.8 | -441007 |
| TA-13 | TA | Tarangire NP | 177523 | -444401 |
| TA-14 | TA | Tarangire NP | 177595.1 | -444307 |
| TA-15 | TA | Tarangire NP | 180261.1 | -447310 |
| TA-16 | TA | Tarangire NP | 180260.7 | -446981 |
| TB-1 | TB | Tarangire NP | 183902.2 | -453155 |
| TB-2 | TB | Tarangire NP | 183768.1 | -453218 |
| TB-3 | TB | Tarangire NP | 182148.9 | -452704 |
| TB-4 | TB | Tarangire NP | 182112.8 | -452564 |
| TB-5 | TB | Tarangire NP | 179855.6 | -452141 |
| TB-6 | TB | Tarangire NP | 179708.8 | -452127 |
| TB-7 | TB | Tarangire NP | 177981.8 | -450880 |
| TB-8 | TB | Tarangire NP | 177902.2 | -450792 |
| TB-9 | TB | Tarangire NP | 176262.9 | -449731 |
| TB-10 | TB | Tarangire NP | 176173.9 | -449653 |
| TB-11 | TB | Tarangire NP | 176600 | -446489 |
| TB-12 | TB | Tarangire NP | 176705.1 | -446403 |
| TB-13 | TB | Tarangire NP | 174817.8 | -444388 |
| TB-14 | TB | Tarangire NP | 174923.5 | -444350 |
| TB-15 | TB | Tarangire NP | 177013.3 | -438641 |
| TB-16 | TB | Tarangire NP | 177137.6 | -438599 |
| TB-17 | TB | Tarangire NP | 175713.3 | -436707 |
| TB-18 | TB | Tarangire NP | 175797.3 | -433914 |
| TB-19 | TB | Tarangire NP | 175752.6 | -433802 |
| TC-1 | TC | Tarangire NP | 173251 | -421652 |
| TC-2 | TC | Tarangire NP | 173220.6 | -421768 |
| TC-3 | TC | Tarangire NP | 174119.1 | -424909 |
| TC-4 | TC | Tarangire NP | 174101.6 | -425026 |
| TC-5 | TC | Tarangire NP | 174093.3 | -427756 |
| TC-6 | TC | Tarangire NP | 174194 | -427822 |
| TC-7 | TC | Tarangire NP | 176538.1 | -429781 |
| TC-8 | TC | Tarangire NP | 176646.3 | -429786 |
| TC-9 | TC | Tarangire NP | 179625.5 | -429866 |
| TC-10 | TC | Tarangire NP | 179737.1 | -429824 |
| TC-11 | TC | Tarangire NP | 181196.1 | -430670 |
| TC-12 | TC | Tarangire NP | 181276.2 | -430743 |
| TC-13 | TC | Tarangire NP | 185134.2 | -432707 |
| TC-14 | TC | Tarangire NP | 185208.6 | -432647 |
| TC-15 | TC | Tarangire NP | 185478.8 | -430603 |
| TC-16 | TC | Tarangire NP | 185353.4 | -430546 |
| BA-1 | BA | Lobosoiret | 196443.2 | -474089 |
| BA-2 | BA | Lobosoiret | 196540.9 | -474048 |
| BA-3 | BA | Lobosoiret | 198862.3 | -475777 |
| BA-4 | BA | Lobosoiret | 198961.9 | -475856 |
| BA-5 | BA | Lobosoiret | 201891.1 | -476718 |
| BA-6 | BA | Lobosoiret | 201973.1 | -476656 |
| BA-7 | BA | Lobosoiret | 204397.6 | -476615 |
| BA-8 | BA | Lobosoiret | 204495.8 | -476564 |
| BA-9 | BA | Lobosoiret | 206114.6 | -475525 |
| BA-10 | BA | Lobosoiret | 206145.5 | -475429 |
| BA-11 | BA | Lobosoiret | 201296 | -496248 |
| BA-12 | BA | Lobosoiret | 201393.9 | -496203 |
| BA-13 | BA | Lobosoiret | 204968 | -492717 |
| BA-14 | BA | Lobosoiret | 205045.9 | -492630 |
| BA-15 | BA | Lobosoiret | 206585.7 | -489703 |
| BA-16 | BA | Lobosoiret | 206612.1 | -489587 |
| BA-17 | BA | Lobosoiret | 207077 | -487013 |
| BA-18 | BA | Lobosoiret | 207040.6 | -486886 |
| BA-19 | BA | Lobosoiret | 207231.9 | -482912 |
| BA-20 | BA | Lobosoiret | 207254.5 | -482776 |
| BB-1 | BB | Kimotorok | 215096.5 | -441900 |
| BB-2 | BB | Kimotorok | 215074.3 | -442017 |
| BB-3 | BB | Kimotorok | 214548.5 | -446742 |
| BB-4 | BB | Kimotorok | 214556.6 | -446849 |
| BB-5 | BB | Kimotorok | 215311.5 | -451567 |
| BB-6 | BB | Kimotorok | 215371.5 | -451687 |
| BB-7 | BB | Kimotorok | 216877 | -455466 |
| BB-8 | BB | Kimotorok | 216897 | -455571 |
| BB-9 | BB | Kimotorok | 218574.7 | -459906 |
| BB-10 | BB | Kimotorok | 218591.7 | -460021 |
| BB-11 | BB | Kimotorok | 219323.5 | -469269 |
| BB-12 | BB | Kimotorok | 219236.6 | -469328 |
| BB-13 | BB | Kimotorok | 215354 | -470747 |
| BB-14 | BB | Kimotorok | 215238.1 | -470780 |
| BB-15 | BB | Kimotorok | 209051.2 | -474730 |
| BB-16 | BB | Kimotorok | 209059.9 | -474853 |
| BB-17 | BB | Kimotorok | 208837.6 | -478110 |
| BB-18 | BB | Kimotorok | 208833.1 | -478251 |

**S3: A table displaying transect, region and coordinates for each Nzi trap set as part of the study**. Tarangire NP = Tarangire National Park.
