## Supplementary 4 for "Caught in a trap: DNA contamination in tsetse xenomonitoring can lead to over-estimates of *Trypanosoma brucei* infection"

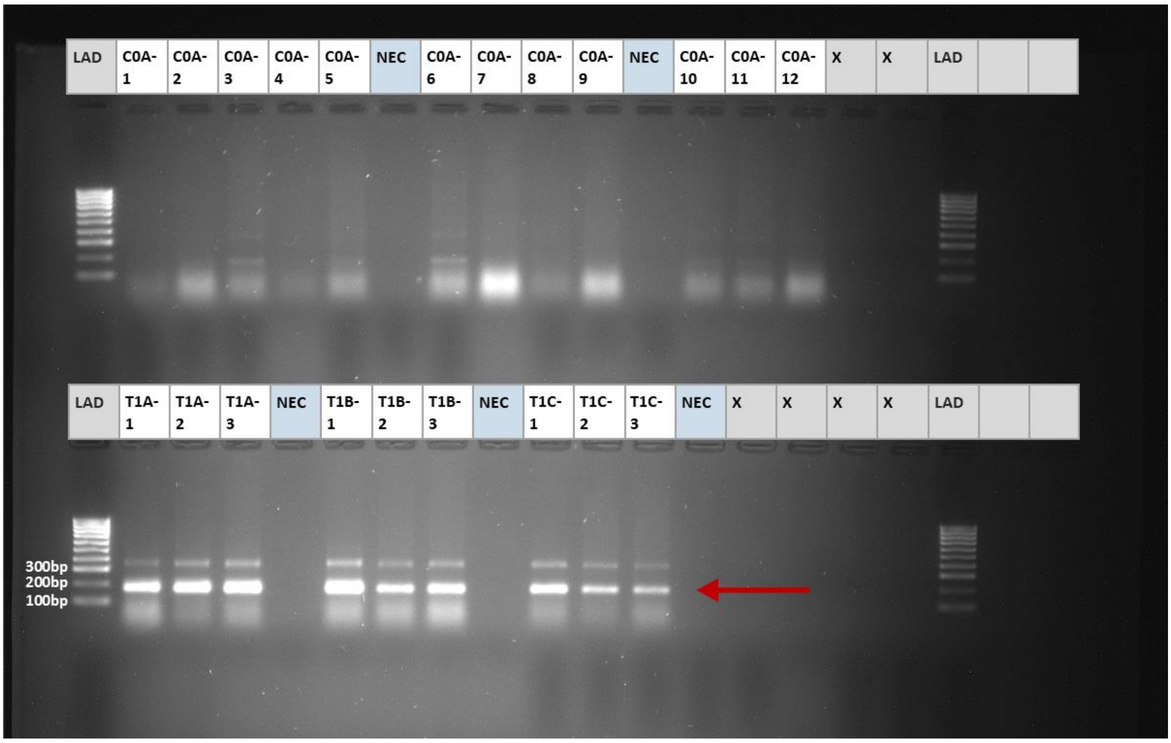


**S4: Gel electrophoresis image from TBR-PCR screening of UFs in C0-A control trap (top row) and naïve flies in T1 control traps (bottom row).** Red arrow indicates target 173bp TBR product. NEC = negative extraction control. LAD = 100bp ladder.
