## Supplementary 5 for "Caught in a trap: DNA contamination in tsetse xenomonitoring can lead to over-estimates of *Trypanosoma brucei* infection"

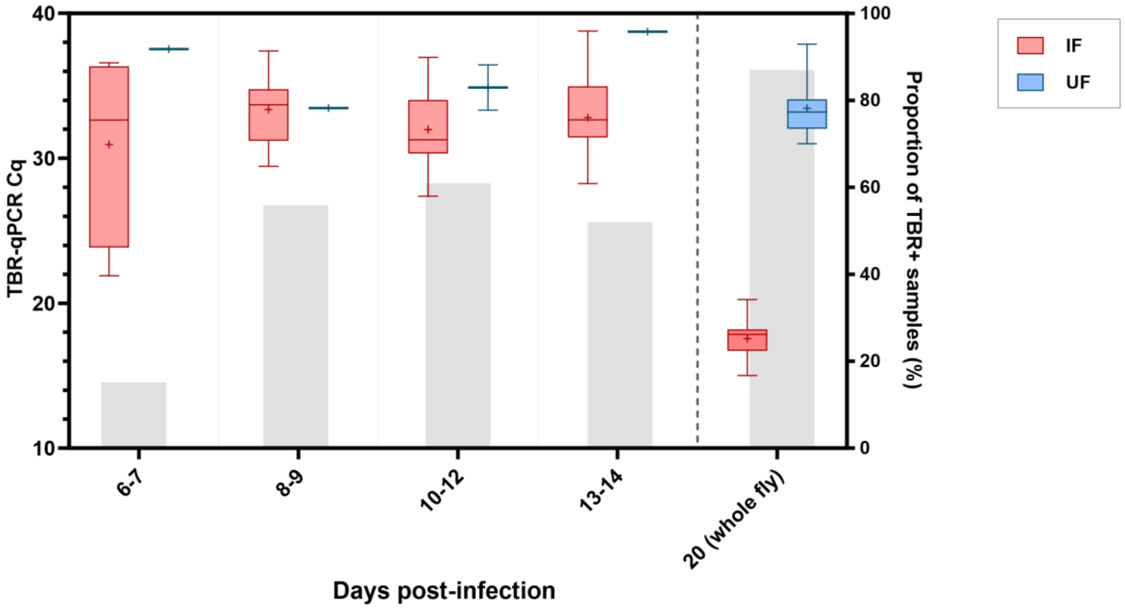


**S5A:** A box-and-whisker plot (left axis) showing Cq values obtained from TBR-qPCR screening of faecal samples at four timepoints and eventual whole fly DNA from a subset of infected (IF) and refractory uninfected flies (UF) that were subject to dissection ante-mortem (n=44). Samples from infected flies are in red, samples from refractory (uninfected) flies are in blue. The bars (right axis) shows the proportion of faecal samples recording TBR-qPCR amplification (where samples were available). The crosses represent the mean Cq values. The amount of *T. brucei* DNA detected in IF samples was consistently higher than that detected in UFs. Where amplification was recorded, there was a significant difference between mean TBR-qPCR Cq values from infected (mean=17.57) and uninfected whole flies at 20 days (mean=33.54, *p*=<0.0001). The midgut infection rate of this subset was 57% (25/44).

| | Infected_Y | Infected_N | | |
|---|---|---|---|---|
| **qPCR_Y** | 40 | 4 | **91%** | *PPV* |
| **qPCR_N** | 5 | 29 | **85%** | *NPV* |
| | **89%** | **88%** | | |
| | *Sensitivity* | *Specificity* | | |

**S5B:** Diagnostic positive predictive value (PPV) and negative predictive value (NPV) calculations for TBR-qPCR screening of tsetse faecal samples as a diagnosis of infection. Faecal samples collected 10-14 days post-inoculation that tested positive (TBR-qPCR) were highly likely to originate from an infected fly, with diagnostic positive predictive value (PPV) of 91% and negative predictive value (NPV) of 85% A positive TBR-qPCR result (‘qPCR_Y’) was any sample recording amplification (Cq < 40). A negative TBR-qPCR result (‘TBR-qPCR_N’) was any sample that did not record amplification. Infected (‘Infected_Y’) was any fly confirmed as having mature midgut infection by microscopy, whilst uninfected (‘Infected_N’) was any fly confirmed as having no visible trypanosome infection by microscopy. Calculations are based on samples collected 10-12 days post-inoculation and/or 13-14 days post-inoculation.
