## Supplementary 6 for "Caught in a trap: DNA contamination in tsetse xenomonitoring can lead to over-estimates of *Trypanosoma brucei* infection"

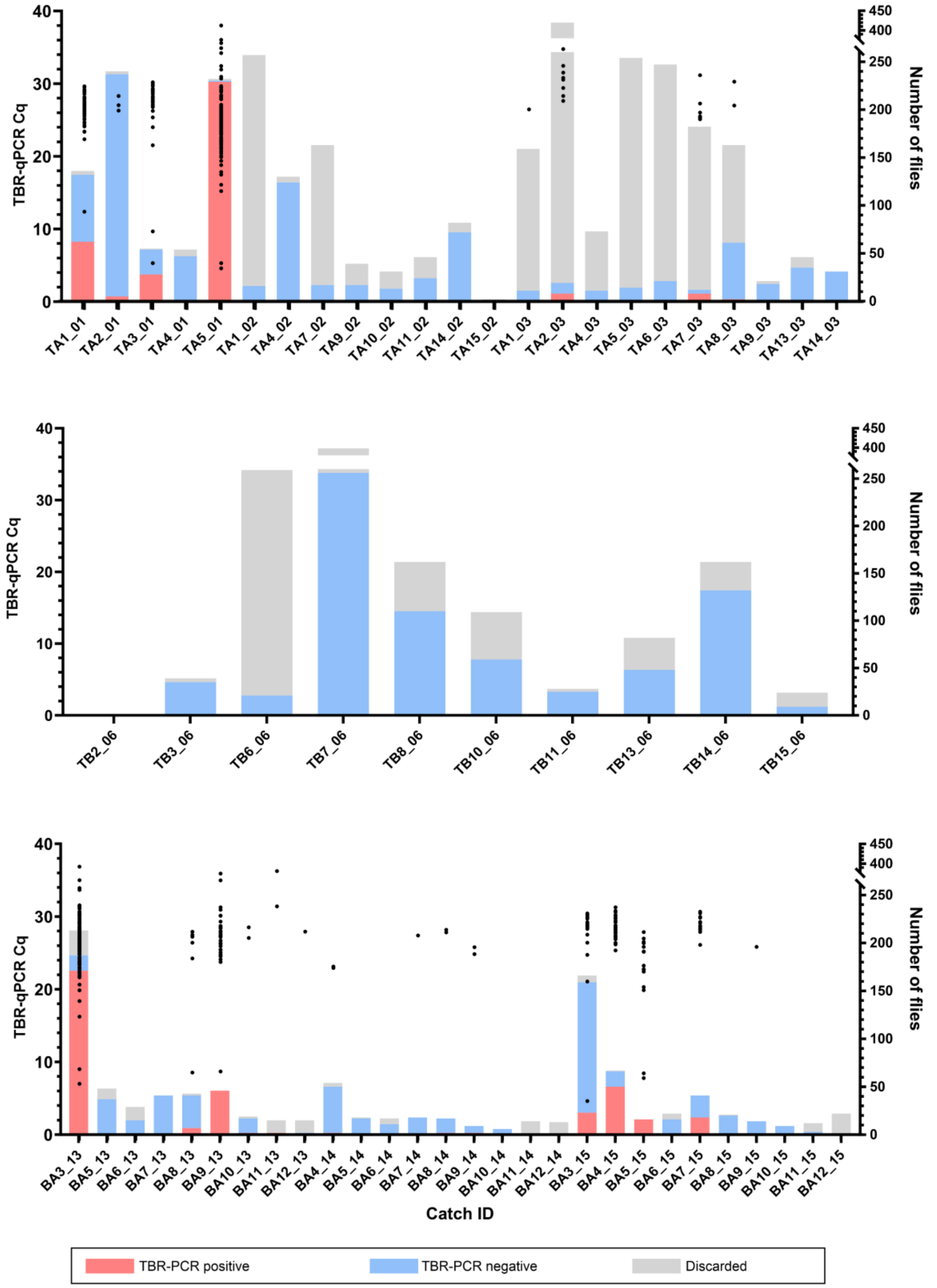


**S6: Plots displaying total catch counts and respective sample TBR-qPCR Cq values for transects TA, TB and BA*.** The left Y axis displays individual fly TBR-qPCR Cq values, plotted as black, circular symbols. The right Y axis displays number of flies caught in each catch, displayed as a stacked bar chart. Red shows the number of flies testing TBR-positive, blue shows the number of flies testing TBR negative, and grey shows the number of flies that were discarded and not collected. *Transect BB is not featured, as it consisted of 1 TBR-negative fly caught in 1 trap (BB17_15).
